# Estrogen withdrawal alters oxytocin signaling in the paraventricular hypothalamus and dorsal raphe nucleus to increase postpartum anxiety

**DOI:** 10.1101/2020.06.16.154492

**Authors:** Valerie L. Hedges, Elizabeth C. Heaton, Claudia Amaral, Lauren E. Benedetto, Clio L. Bodie, Breanna I. D’Antonio, Dayana R. Davila Portillo, Rachel H. Lee, M. Taylor Levine, Emily C. O’Sullivan, Natalie P. Pisch, Shantal Taveras, Hannah R. Wild, Amy P. Ross, H. Elliott Albers, Laura E. Been

## Abstract

**Background:** Estrogen increases dramatically during pregnancy, but quickly drops below pre-pregnancy levels at birth and remains suppressed during the postpartum period. Clinical and rodent work suggests that this postpartum drop in estrogen results in an “estrogen withdrawal” state that is related to changes in affect, mood, and behavior. Most studies examining the effect of estrogen withdrawal on the brain have focused solely on the hippocampus.

**Methods:** We used a hormone-simulated pseudopregnancy model in Syrian hamsters, a first for this species. Ovariectomized females were given daily injections to approximate hormone levels during gestation and then withdrawn from estrogen to simulate postpartum estrogen withdrawal. Subjects were tested for behavioral assays of anxiety and anhedonia during estrogen withdrawal. Following sacrifice, neuroplasticity in oxytocin-producing neurons in the paraventricular nucleus of the hypothalamus (PVH) and its efferent targets was measured.

**Results:** Estrogen-withdrawn females had increased anxiety-like behaviors in the elevated plus and open field, but did not differ from controls in sucrose preference. Furthermore, estrogen-withdrawn females had more oxytocin-immunoreactive cells and oxytocin mRNA in the PVH, as well as an increase in oxytocin receptor density in the dorsal raphe nucleus (DRN). Finally, blocking oxytocin receptors in the DRN during estrogen withdrawal prevented the high-anxiety behavioral phenotype in estrogen-withdrawn females.

**Conclusions:** Estrogen withdrawal alters oxytocin signaling in the PVH and DRN to increase anxiety-like behavior during the postpartum period. More broadly, these experiments suggest Syrian hamsters as a novel organism in which to model the effects of postpartum estrogen withdrawal on the brain and anxiety-like behavior.

## Introduction

During pregnancy and the postpartum period, many hormones (e.g., estrogens, progesterone, beta-endorphins, oxytocin, corticosterones) fluctuate and likely contribute uniquely to peripartum changes in physiology, behavior, and mood/affect. Among these hormones, peripartum fluctuations in estrogen levels are particularly dramatic: estrogen, synthesized primarily in the placenta, rises 100-1000-fold during the third trimester (1). Following birth and the expulsion of the placenta, however, estrogen levels quickly drop to below pre-partum levels and remain suppressed until ovulation resumes weeks to months later (2). It has been hypothesized that the rapid and dramatic drop in estrogen during the postpartum period results in an “estrogen withdrawal state” that contributes to the etiology of postpartum psychological disorders (3). In support of this idea, several studies have found that symptoms of postpartum depression can be attenuated by estradiol treatment in women (4–6).

How estrogen withdrawal contributes to the etiology of peripartum psychological disorders is unclear. To better understand the impact of this dramatic endocrine change, some researchers have turned to rodent models. In particular, several groups have employed a hormone-simulated pseudopregnancy (HSP) model to directly test the impact of estrogen withdrawal on the brain and behavior. Initially developed in female Long-Evans rats (7), this model has successfully been replicated (8) and extended to Sprague-Dawley rats (9–13), and ICR mice (14,15), suggesting it is a highly reproducible way to model the impact of postpartum estrogen withdrawal on behavior across multiple species. Consistent with the clinical literature, estrogen withdrawal following HSP results in a behavioral phenotype that may reflect depressed mood (7–12) and/or heightened anxiety (11,14,15) in rodents. In many cases, these behavioral changes can be prevented by continued administration of estradiol during the postpartum period, also matching the data from human studies.

An extreme, systemic endocrine shift like postpartum estrogen withdrawal is likely to impact multiple, distributed brain regions that have distinct effects on behavior. Despite this, the majority of studies that have used HSP to examine how postpartum estrogen withdrawal impacts the brain have focused on the hippocampus (8,11,13,14). Only one study to date has examined the impact of postpartum estrogen withdrawal outside of the hippocampus. In an elegant series of experiments, Yang et al. (2017) demonstrated that estrogen withdrawal following HSP in mice led to an increase in anxiety-like behavior, and that this behavior change was mediated in part by G-protein coupled estrogen receptor 1-mediated inhibition of LTD in the basolateral amygdala. It is clear then that extra-hippocampal brain regions are impacted by postpartum estrogen withdrawal, and that these neuroplastic changes may underlie different aspects of postpartum behavioral changes.

The paraventricular hypothalamus (PVH) is a highly-conserved subnucleus of the hypothalamus, which secretes neuropeptides either into peripheral circulation via the neurohypophysial system, or to neural targets via centrally-projecting neurons (16). In particular, oxytocin (OT), synthesized primarily in the PVH and supraoptic nucleus (SON), is considered a key modulator of physiology, behavior, and affective processes associated with pregnancy and the postpartum period (17). Oxytocin receptors (OTR) are located in several estrogen-sensitive brain nuclei and are functionally implicated in maternal behaviors, anxiety, depression, social cognition, and sexual behavior (18,19). Furthermore, estrogen increases both the secretion of OT and the expression of OTRs in the brain (20), suggesting the oxytocinergic system as a likely target for neuroplasticity underlying behavioral changes during postpartum estrogen withdrawal. Using HSP in Syrian hamsters, a well-established rodent model of reproductive and affective behaviors, we investigated the role of postpartum estrogen withdrawal on functional neuroplasticity in the PVH and its efferent targets.

## Methods and Materials

### Subjects

For all experiments, adult female Syrian hamsters were purchased from Charles River Laboratories (Wilmington, Massachusetts) at approximately 55 days of age. Female hamsters were housed individually in polycarbonate cages (19” x 10.5” x 8”) with aspen bedding. Hamsters are solitary, territorial animals and individual housing is standard and non-stressful (21,22). Animal holding rooms were temperature- and humidity-controlled and maintained on a reversed 14-hr light/10-hr dark period. Food and water were provided *ad libitum*. All animal procedures were carried out in accordance with the National Institutes of Health Guide for the Care and Use of Laboratory Animals and approved by the Institutional Animal Care and Use Committee. The experimental groups and numbers of subjects for each experiment can be found in Table 1.

**Table 1.** Experimental groups, numbers of animals, and dependent variables for each experiment.

| Experiment | Groups (n) | Dependent Variables |
| --- | --- | --- |
| 1A: Effect of Estrogen Withdrawal on Anxiety-Like Behaviors | Withdrawn (n = 8)<br>Sustained (n = 8)<br>No Hormone (n = 8) | Elevated Plus<br>Open Field<br>Locomotor behavior |
| 1B: Effect of Estrogen-Withdrawal on Anhedonia | Withdrawn (n = 8)<br>Sustained (n = 8) | Sucrose Preference<br>Sucrose Consumed |
| 2A: Effect of Estrogen-Withdrawal on Oxytocin-Producing Cells | Withdrawn (n = 8)<br>Sustained (n = 8)<br>No Hormone (n = 8) | Number of OT-ir cells in PVH and SON and NeuN-ir cells in PVH |
| 2B: Effect of Estrogen-Withdrawal on OT Gene Expression in PVH | Withdrawn (n = 6)<br>Sustained (n = 6) | OT mRNA (qPCR) in PVH |
| 3: Effect of Estrogen-Withdrawal on OTR density in PVH efferents | Withdrawn (n = 8)<br>Sustained (n = 8)<br>No Hormone (n = 8) | OTR density in MeA, BNST, NAc Core, NAc Shell, and Dorsal Raphe |
| 4: Effect of Blocking OTRs in DRN during Estrogen-Withdrawal on Anxiety-Like Behaviors | Withdrawn VEH (n = 8)<br>Sustained VEH (n = 8)<br>Withdrawn OTA (n = 9)<br>Sustained OTA (n = 7) | Elevated Plus<br>Open Field |

### Hormone-Simulated Pseudopregnancy (HSP)

Following recovery from ovariectomy (see Surgery), the HSP protocol (7) was initiated. In this model, ovariectomized females are given daily injections of estrogen and/or progesterone in doses that are known to reliably induce maternal behaviors in nulliparous ovariectomized rats (23–25). For these experiments, we have modified the protocol to fit the shorter gestational timeline (17d) of Syrian hamsters (26).

All hamsters were administered daily subcutaneous injections of hormone dissolved in cottonseed oil vehicle (Sigma, St. Louis, MO, USA) over a 22-day period. On days 1-12, hamsters received a low dose (2.5 μg) of estradiol benzoate (Sigma) and a high dose (4 mg) of progesterone (Sigma). On days 13-17, hamsters received a high dose of estradiol benzoate (50 μg). On days 18-22, hamsters were divided into two experimental conditions: estrogen-withdrawn (“withdrawn”) females received daily vehicle injections, modeling the dramatic drop in estrogen that occurs during the postpartum period, while females in the estrogen-sustained (“sustained”) group continued to receive daily injections of a high dose (50 μg) of estradiol. In some experiments, both withdrawn and sustained females were compared to a “no hormone” condition, in which females received daily vehicle (oil) injections for all 22 days of the hormone simulated pregnancy.

### Behavior Testing

#### Elevated Plus

An elevated plus test was used to assess anxiety-like behavior. This test was chosen because it has previously been used to measure anxiety-like behavior following HSP in rats and mice (11,14,15). The apparatus consists of a plus-shaped maze elevated 73 cm above the floor, with two open arms (51 × 11.5 cm) and two enclosed arms (51 × 11.5 × 39.5 cm). On day 20 or 21, animals were placed in the center of the maze facing a closed arm and allowed to freely explore the maze for five min. Tests were recorded and the amount of time spent in each arm, as well as the total distance traveled and average velocity were quantified using Noldus Ethovision XT (version 14, Noldus Information Technology, Wageningen, The Netherlands).

#### Open Field

An open field test was used to measure spontaneous locomotor activity. This test was chosen because it has been used previously following HSP in rats and mice (11,14,15). The apparatus consists of an open field (40.5 × 40.5 × 30cm) composed entirely of solid, light grey plastic (Maze Engineers, Glenview, IL, USA). On day 20 or 21, animals were placed in the center of the apparatus and allowed to freely explore for five min. Tests were recorded and the amount of time spent in the center of the field, periphery of the field, as well as the total distance traveled and average velocity were quantified using Noldus Ethovision XT.

#### Sucrose Preference

*Test*. A sucrose preference test was used to assess anhedonia, a core feature of depressed mood. This test was chosen because sucrose consumption and preference has previously been shown to decrease following HSP in rats (8). Further, other common assays of depressed mood in rodents (e.g., forced swim, tail suspension) are not possible in hamsters.

Hamsters were given a single bottle of a 3% sucrose solution for 48 h in their home cage to allow for acclimation to the solution. Subsequently, hamsters were presented with two bottles: one bottle filled with tap water and one with 3% sucrose solution for 48 h to acclimate them to the presence of two bottles in their cage. Fluid intake, preference data, and body weight data were collected during 1 h tests six times over two weeks to establish baseline consumption. Fluid intake was calculated by measuring bottle weight prior to, and immediately after, each one-hour exposure. Following the establishment of a baseline, a 1-hour two-bottle choice test was performed during “early pregnancy” (Day 12), “late pregnancy” (day 17) and “postpartum” days 2 and 3 (days 20 and 21) of the hormone simulated pregnancy. All two-bottle choice tests were done in the first hour of the dark cycle, and bottle position was alternated to control for side preference. After each test, bottles were removed and replaced with regular water bottles.

### Surgery

All surgery was conducted using aseptic surgical technique and under isoflurane anesthesia (2-3% vaporized in oxygen, Piramal, Bethlehem, PA, USA). Analgesic (Meloxicam, 2mg/kg, Portland, ME, USA) was administered subcutaneously immediately prior to the start of surgery and for three days postoperatively.

#### Ovariectomy

In order to remove the endogenous source of circulating estrogen and progesterone, females were bilaterally ovariectomized prior to the initiation of the hormone simulated pregnancy. Briefly, subjects’ bilateral flanks were shaved and cleaned with three alternating scrubs of 70% ethanol and Betadyne before being transferred to a sterile surgical field. Anesthesia was maintained via a nosecone. Subjects’ ovaries were extracted via bilateral flank incisions and removed via cauterization of the uterine horn and blood vessels. Polydioxanone absorbable suture (Ethicon, Sommerville, NJ, USA) and wound clips (Fine Science Tools, Foster City, CA, USA) were used to close the smooth muscle and skin incisions, respectively.

#### Dorsal Raphe Cannulation

At least 1 week following ovariectomy, subjects in Experiment 4 were stereotaxically implanted with 26-guage stainless steel guide cannulae (Plastics One, Roanoke, VA, USA) targeting the dorsal raphe nucleus. Briefly, the dorsal scalp was shaved and subjects were secured with ear bars in a stereotaxic apparatus such that their skull was level in the anterior-posterior (A-P) and medial-lateral (M-L) planes. Anesthesia was maintained via a nosecone and the scalp was cleaned with three alternating scrubs of 70% ethanol and Betadine. Following a midline scalp incision, the skin and temporal muscles were retracted to expose the skull and A-P and M-L stereotaxic coordinates for the dorsal raphe (27) were measured relative to bregma (A-P: -4.5mm, M-L: ∓1.8mm). A hand-operated drill was used to drill three holes in the skull: one for the guide cannula and two for stainless steel bone screws (Plastics One). Following placement of the bone screws, the guide cannula was lowered 2.3 mm below dura using a stereotaxic arm (David Kopf Instruments, Tujunga, CA, USA). Once implanted, the cannula was fixed to the skull using dental acrylic (Patterson Dental, St. Paul, MN, USA) and a stainless-steel dummy cannula (Plastics One) was inserted into the cannula shaft to prevent occlusion.

### Drug Injections

Oxytocin receptor antagonist (OTA) was diluted in sterile saline and stored in aliquots of 10 μL at −20°C until use. On behavior testing days (days 20 and 21), subjects with DRN cannulae were injected with either 250 nL of OTA (900 µM, provided by Maurice Manning to Elliott Albers) or 250 nL of sterile saline 10 min before the test. This dose has been demonstrated to produce behavioral effects in female Syrian hamsters (28,29). Drug delivery was accomplished using an automated syringe pump (New Era Pump Systems, Inc, East Farmingdale, NY, USA) and injection needles extended 3 mm below the guide cannula (Plastics One). Following injection, the injection needle was left in the brain for an additional 30 sec, to allow the injected solutions to diffuse away from the needle. After removing the injection needle, dummy stylets were replaced, and hamsters were then transferred back to their cages until testing.

### Histology and Tissue Processing

#### Perfusion and Immunohistochemistry

For experiments requiring immunolocalization of oxytocin or Neuronal Nuclei (NeuN) protein, subjects were sacrificed by intracardial perfusion. On day 22, subjects were given an overdose of sodium pentobarbital (Beuthanasia-D Special, 22 mg/100g body weight, Merck Animal Health, Madison, NJ, USA). Once at surgical plane, subjects were transcardially perfused with approximately 200 mL of 25 mM PBS (pH 7.4), followed by 200 mL of 4% paraformaldehyde (Electron Microscopy Sciences, Hatfield, PA, USA). Brains were immediately removed and post-fixed in 4% paraformaldehyde overnight (4° C) and then cryoprotected for 48 hr in 30% sucrose in PBS. Coronal sections (40-μm) of brain tissue were sectioned on a cryostat (−20°C), collected in a 1:6 series, and stored in cryoprotectant until immunohistochemical processing.

Procedures for oxytocin and NeuN immunohistochemistry were identical except for the specific primary and secondary antibodies used. Sections were removed from cryoprotectant and rinsed 5 × 5 min in 25 mM PBS. To reduce endogenous peroxidase activity, tissue sections were incubated in 0.3% hydrogen peroxide for 15 min. After 5 × 5 min rinses in PBS, sections were incubated in a mouse monoclonal primary antibody against oxytocin (1:10,000, MAB5296, Millipore, Burlington, MA, USA) or rabbit polyclonal antibody against NeuN (1:20,000, ABN78, Millipore) in 0.4% Triton-X 100 for 24 hr at room temperature. After incubation in primary antibody, sections were rinsed in PBS and then incubated for 1 hr in the appropriate biotinylated secondary antibody (anti-mouse or anti rabbit, 1:600 Jackson ImmunoResearch, West Grove, PA, USA) in PBS with 0.4% Triton-X 100. Sections were rinsed again in PBS and then incubated for 1 hr in avidin-biotin complex (4.5 μL each of A and B reagents/ml PBS with 0.4% Triton-X 100, ABC Elite Kit, Vector Laboratories, Burlingame, CA, USA). After rinsing in PBS, sections were incubated in 3,3’-diaminobenzidine HCl (0.2 mg/mL, Sigma) and hydrogen peroxide (0.83 μL/mL, Sigma) for 10 min, yielding an orange-brown product. The reaction was stopped by rinsing sections in PBS. For both OT and NeuN, a subset of sections was run through the immunohistochemistry protocol with the primary antibody omitted. Stained tissue sections were mounted onto subbed glass slides and allowed to air-dry overnight. Slides were then dehydrated in alcohols, cleared in xylenes, and cover slipped using Permount (Fisher Scientific, Waltham, MA, USA).

A light microscope (Nikon Eclipse E200) using a color camera (Diagnostic Instruments, Sterling Heights, MI, USA) and SpotBasic software was used to acquire images. For OT-ir sections, 10X images of the bilateral PVH and the unilateral SON (2 unilateral images per animal) were acquired. For NeuN-ir sections, unilateral images of the PVH were acquired at 40X magnification (2 unilateral images per animal) All cell counts were done by experimenters blind to the condition of the subject.

#### Receptor Autoradiography

For experiments measuring changes in oxytocin receptor density following a hormone simulated pregnancy, subjects were euthanized via rapid decapitation. On day 22, subjects were given an overdose of sodium pentobarbital (Beuthanasia-D Special, Merck Animal Health), and their brains were rapidly extracted, flash frozen, and transported on dry ice to Georgia State University where they were sectioned (40 µm) on a cryostat (−20°C) and mounted on Superfrost Plus slides (Fisher Scientific) for oxytocin receptor autoradiography.

Oxytocin receptor binding was determined with the I^125^-labeled ornithine vasotocin analog Vasotocin, d(CH_2_)_5_[Tyr(Me)^2^,Thr^4^,Orn^8^,[^125^I]Tyr9-NH2] (Perkin-Elmer, Waltham, MA, USA). The tissue was thawed, dried, and then fixed in 0.1% paraformaldehyde for 2 min. The slides were then rinsed twice for 10 min each in buffer (50 mM Tris, pH 7.4) and were then incubated in tracer buffer (0.35 mM bacitracin, Sigma-Aldrich; 0.015 mM bovine serum albumin, Sigma-Aldrich; 100 nM I^125^ vasotocin analog) for 1 hr. The slides were then rinsed twice for 5 min each and then for 35 min with agitation in buffer (50 mM Tris, 21 mM MgCl). All incubations and washes were performed at room temperature. Finally, the slides were dipped in 4°C deionized water and allowed to dry. The slides and a C^14^ standard calibration strip (American Radiolabeled Chemicals, St. Louis, MO, USA) were loaded into autoradiography cassettes and exposed to film (Kodak, Rochester, NY, USA) for 7 days at room temperature.

Densitometry analysis was performed on five regions of interest: the medial amygdala (MeA), bed nucleus of the stria terminalis (BNST), Nucleus Accumbens Core (NAcC), Nucleus Accumbens Shell (NAcSh) and Dorsal Raphe Nucleus (DRN). Analyses were performed by an individual blind to the experimental condition of each subject using Scion Image software (NIH, Bethesda, MD, USA) and a lightbox (Imaging Research, Inc., Ontario, Canada) attached to a camera (Panasonic, Newark, NJ, USA). Standard curves were created using the C^14^ microscales on the standard calibration strip. For each brain area of interest, three tissue sections located 60 μm apart were analyzed on the right and left sides of the brain, except for dorsal raphe nuclei sections, which were analyzed along the midline of the brain. With the exception of the MeA, a 0.35 mm^2^ box was placed over the center of each brain area, and the optical density was recorded. A 0.35 mm x 0.75 mm box was used to analyze the MeA in order to measure dorsal and ventral MeA simultaneously. Background binding was subtracted from this measurement. Optical densities were calculated as disintegrating units per min per mg tissue (dpm/mg).

#### Cannulae Placement Verification

At the conclusion of the DRN cannulation experiment, subjects were given an overdose of sodium pentobarbital (Beuthanasia-D Special). An injection needle was lowered into the cannula and 250 nL of India Ink (Dr. Ph. Martin’s, Salis International, Oceanside, CA, USA) was infused to mark the location of the injection site using an automated syringe pump. Subjects were then decapitated and their brains were removed and post-fixed in paraformaldehyde for 48 h. A cryostat (Leica CM1850, Leica Biosystems, Buffalo Grove, IL, USA) was used to collect 40-µm sections through the rostral-caudal extent of the dorsal raphe. Sections were mounted onto slides and a brightfield microscope (Nikon) using SpotBasic software (Diagnostic Instruments) was used to acquire an image of the location of the ink injection. Only subjects whose ink injection placement was inside the dorsal raphe were included in the analysis.

#### qPCR

Tissue samples were homogenized and RNA was extracted using an RNAeasy Mini Kit (Quiagen, Hilden, Germany) and reverse transcribed using a Transcriptor First-Strand cDNA Synthesis kit (Roche Diagnostics, Indianapolis, Indiana, USA). Primers for Oxytocin were designed using known sequences from the Syrian hamster (Forward: 5′-CGG-CCT-GCT-ACA-TCC-AGA-AT-3′, Reverse: 5′-ATG-ACC-CTC-TTG-GCT-CCA-C-3′ GenBank Accession HM357357.1). qPCR amplifications were performed on a StepOne Plus Real-Time PCR System (Applied Biosystems, Foster City, CA, USA) using SYBR Green (PowerUp SYBR Green Master Mix, Applied Biosystems). PCR for individual cDNA samples were performed in triplicate. Threshold values were calculated using the second derivative max method and standardized to the housekeeping gene Gapdh (primers designed from Syrian hamster sequence: Forward: 5′-GTC-TAC-TGG-CGT-CTT-CAC-CAC-3′, Reverse: 5′-ATG-ACC-CTC-TTG-GCT-CCA-C-3′, GenBank Accession ABD77188.1). The thermal cycling program used a UDG activation step at 50°C for 2 min and a Dual-Lock DNA Polymerase step at 95°C for 2 min, followed by 40 cycles consisting of a 15 sec denaturing step at 95°C, an annealing/extending step for 1 min at 60°C. At the end of each cycling program, a measurement of fluorescent intensity was taken and a melting curve was generated.

### Statistical Analysis

All statistics were run on Prism Software (Graphpad, San Diego, CA, USA). In Experiment 1, difference scores were calculated for the elevated plus (time spent in closed arms - time spent in open arms) and open field (time spent in periphery - time spent in center) and one-way ANOVAs were used to determine whether difference scores varied by experimental condition. Tukey’s HSD post hoc tests were used to examine significant omnibus tests for group differences. Sucrose preference was calculated as the sucrose consumed per 100g of body weight divided by the total liquid consumed (sucrose and water) per 100g of body weight. Unpaired *t*-tests were used to determine whether preference scores or volume of sucrose consumed varied by hormone condition.

In Experiment 1, one-way ANOVAs or unpaired t-tests were used to determine whether anxiety-like behavior or sucrose preference differed across hormone condition. In Experiment 2, one-way ANOVAs or unpaired two-tailed t-tests were used to determine whether cell counts (OT or NeuN) varied by experimental conditions. Tukey’s HSD post hoc tests were used to examine significant omnibus tests for group differences. Likewise, unpaired two-tailed t tests were used to determine whether OT gene expression differed between experimental condition. In Experiment 3, one-way ANOVAs were used to determine whether optical densities (dpm/mg) varied by experimental conditions. Tukey’s HSD post hoc tests were used to examine significant omnibus tests for group differences. In Experiment 4, difference scores were calculated for the elevated plus and open field and two-way ANOVAS were used to probe for interactions and main effects of hormone condition (sustained vs. withdrawn) and drug condition (OTA vs. vehicle). t tests with Bonferroni corrections for multiple comparison were used to explicate interactions.

## Results

### Experiment 1: Effect of estrogen withdrawal on behavioral measures of anxiety and depression

#### Elevated Plus

There was a significant effect of hormone condition on the amount of time spent in the closed arms minus the amount of time spent in the open arms of the elevated plus (F(2, 21) = 14.72, p = .0001). Post hoc tests revealed that sustained females did not differ from no hormone females with regard to where they spent their time in the elevated plus (p = 0.1184); animals in both of these groups spent roughly equivalent amounts of time in both arms. However, withdrawn females differed from both no hormone females (p = 0.0091) and sustained females (p < 0.0001); animals in the withdrawn group spent more time on average in the closed arms than the open arms, indicating a higher anxiety phenotype (Figure 1A).

**Figure 1:**
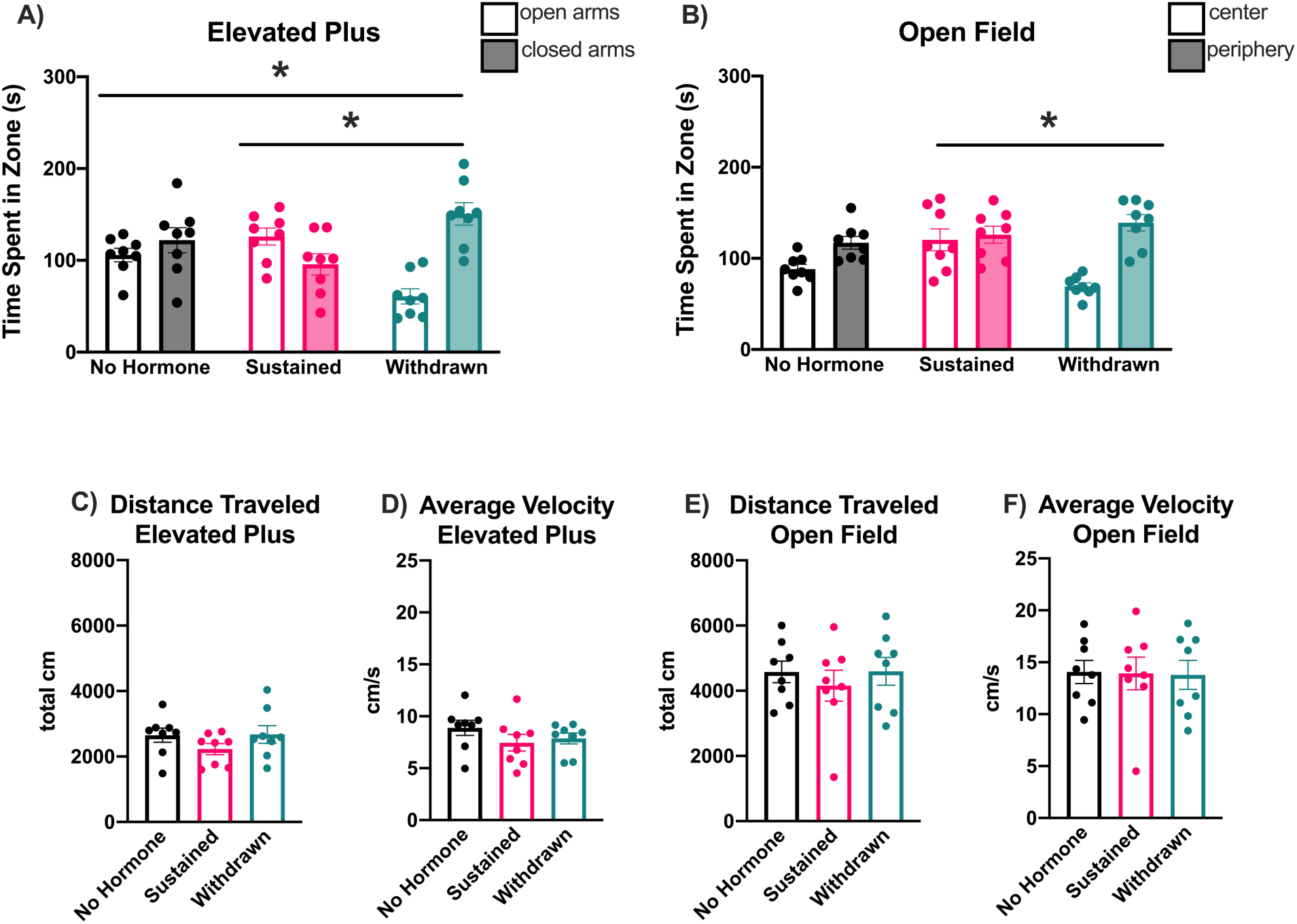
Effect of Estrogen Withdrawal on Anxiety-Like Behaviors. A) Withdrawn females showed a high-anxiety behavioral phenotype in the elevated plus compared to sustained females or no hormone controls. B) Withdrawn females showed a high-anxiety behavioral phenotype in the open field compared to sustained females, but did not differ from no hormone controls. These differences in anxiety-like behavior are not mediated by a more general locomotor deficit, as there is no effect of hormone condition on distance traveled (C and E) or average velocity (D and F) in either test. Data presented as mean ± SEM, * *p* < 0.05.

#### Open Field

There was a significant effect of hormone condition on the amount of time spent in the periphery minus the amount of time spent in the center of the open field (F(2, 21) = 5.457, p = 0.0123). Post hoc tests revealed that no hormone females did not differ from sustained females with regard to where they spent their time in the open field (p = 0.4781). Likewise, no hormone females did not differ from withdrawn females with regard to where they spent their time in the open field (p = 0.1176). However, sustained females differed from withdrawn females (p = 0.0100); animals in the withdrawn groups spent more time on average in the periphery than in the center, indicating a higher anxiety phenotype (Figure 1B).

#### Locomotor Behavior

To assess whether significant effects of hormone condition in the open field and elevated plus could be explained by broader locomotor deficits, distance traveled (total cm) and average velocity (cm/s) were compared across hormone conditions in both apparatuses. In the elevated plus, there was no effect of hormone condition on distance traveled (F(2,21) = 1.279, p = 0.2990, Figure 1C) or average velocity (F(2,21) = 1.134, p = 0.3407, Figure 1D). Likewise, in the open field, there was no effect of hormone condition on distance traveled (F(2,21) = 0.3623, p = 0.7003, Figure 1E) or average velocity (F(2,21) = 0.01055, p = 0.9895, Figure 1F). This suggests that group differences in the elevated plus and open field are reflective of differences in anxiety-like behaviors rather than nonspecific effect of drug or hormone condition on locomotion.

#### Sucrose Preference

There was no effect of hormone condition on sucrose preference during early pregnancy (t(13) = 0.1892, p = 0.8528), late pregnancy (t(14) = 1.682, p = 0.1148), postpartum day 2 (t(14) = 1.238, p = 0.2360), or postpartum day 3 (t(13) = 2.101, p = 0.05650). Similarly, there was no effect of hormone condition on sucrose consumption during early pregnancy (t(13) = 0.2879, p = 0.7779), late pregnancy (t (13) = 1.981, p = .0691), postpartum day 2 (t(13) = 0.9820, p = 0.3440), or postpartum day 3 (t(13) = 0.2371, p = 0.8166). Together, these data suggest that HSP and subsequent estrogen withdrawal do not impact sucrose preference (supplementary Figure 1).

### Experiment 2: Effects of estrogen withdrawal on oxytocin neurons in the PVH

To assess whether OT-producing neurons differed between hormone groups, the number of OT-immunoreactive (-ir) neurons in the PVH (Figure 2A) and supraoptic nucleus (SON, Figure 2B) was counted and compared between groups. There was a significant effect of hormone condition on the number of OT-ir cells in the PVH (F(2,21) = 22.67, p < 0.0001). Post hoc tests revealed that while the no hormone and sustained groups did not differ from each other (p = 0.7191), the withdrawn group had significantly more OT-ir cells than both the no hormone group (p < 0.0001) and the sustained group (p < 0.0001) (Figure 2C). There was no effect of hormone condition on the number of OT-ir cells in the SON, F(2,21) = 0.6547, p = 0.5299) (Figure 2D). Sections that were run with the primary antibody omitted showed no immunostaining.

**Figure 2:**
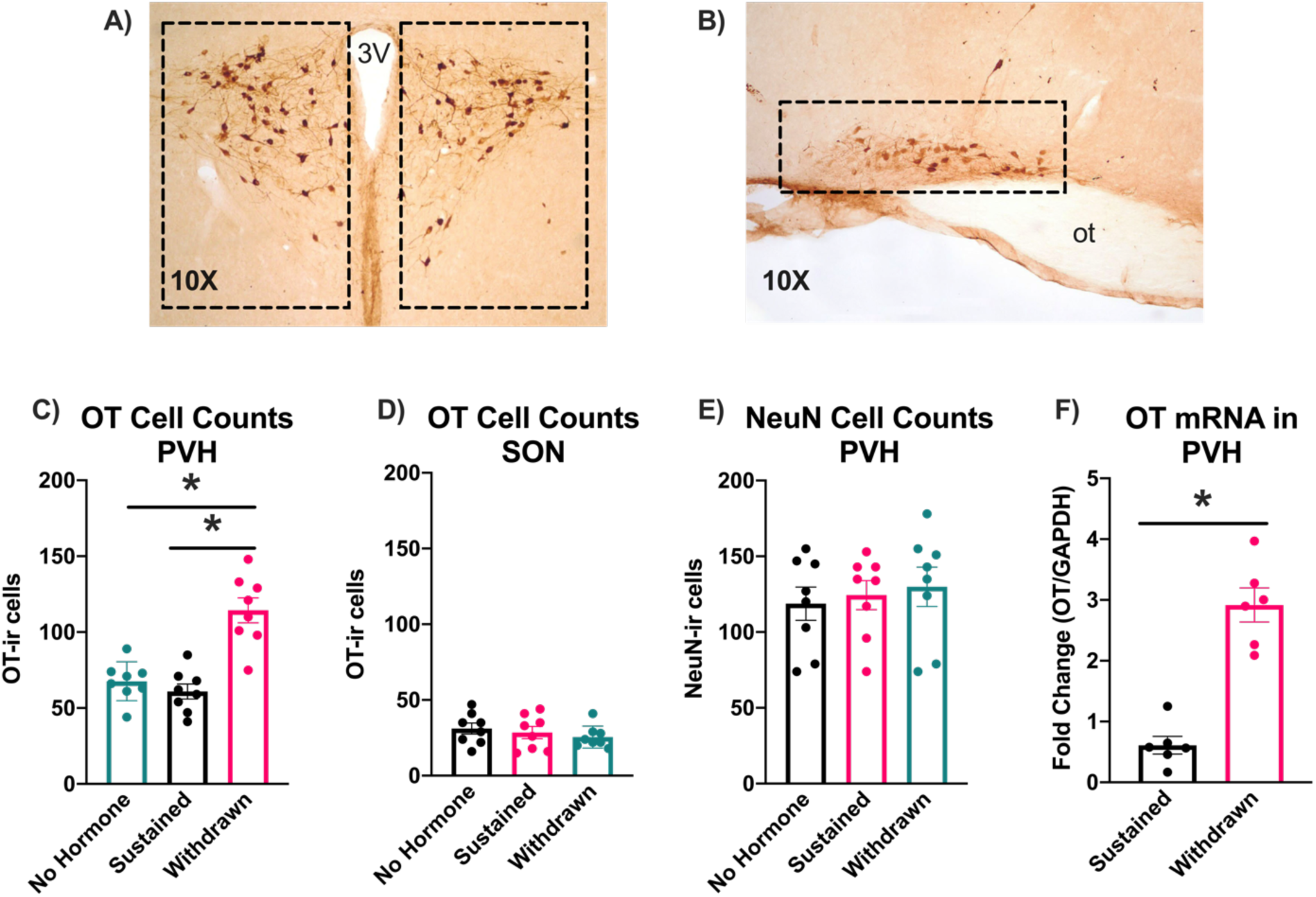
Effect of Estrogen-Withdrawal on Oxytocin-Producing Cells. A) Representative OT-ir staining and counting domains in the PVH. B) Representative OT-ir staining and counting domains in the SON. C) In the PVH, the total number of OT-ir cells was significantly higher in withdrawn females than in sustained or no hormone controls. D) In the SON, the total number of OT-ir cells did not differ between hormone conditions. E) The total number of NeuN-ir cells in the PVH did not differ across hormone conditions. F) There was an approximately three-fold increase in OT gene expression in withdrawn females compared to estrogen sustained females. Data presented as mean ± SEM, * *p* < 0.05.

To assess whether the increase in OT-ir neurons could be explained by an increase in the total number of neurons, the number of NeuN-ir neurons was counted and compared between groups. There was no effect of hormone condition on the total number of NeuN-ir neurons in the PVH (F(2, 21) = 0.2457, p = 0.7843), suggesting that the increase in OT-ir neurons is unlikely to be the result of neurogenesis (Figure 2E).

An additional possibility is that estrogen withdrawal led to a decrease in OT release from PVH cells, causing more OT to build up in the terminal and, in turn, an increase in the detection of immunoreactive cells. In this case, we would not expect OT gene expression to differ between estrogen-sustained and estrogen withdrawn females. To test this, qPCR was used to assess OT mRNA in the PVH of a separate cohort of sustained and withdrawn females. There was a significant increase in OT mRNA in withdrawn animals compared to sustained animals (t(10) = 7.318, p < 0.001), suggesting that the increase in OT-ir cells is unlikely to reflect a decrease in OT release (Figure 2F).

### Experiment 3: Effects of estrogen withdrawal on oxytocin receptors in PVH efferents

To determine whether the increase in oxytocin-ir following estrogen withdrawal was accompanied by a change in oxytocin receptors in PVH efferents, receptor autoradiography was used to measure OTR density in the MeA, BNST, NAc core, NAc shell, and DRN. There was no effect of hormone condition on OTR density in the MeA (F(2,21) = 1.872, p = 0.1785, Figure 3A), BNST (F(2,21) = 1.746, p = 0.1989, Figure 3B), NAc core (F(2,21) = 1.122, p = 0.3409, Figure 3C), or NAc shell (F(2,21) = 1.736, p = 0.2006, Figure 3D). However, there was a significant effect of hormone condition on OTR density in the DRN (F(2,21) = 13.43, p = 0.0002). Post hoc tests revealed that while the no hormone group did not differ from the sustained group (p = 0.4521), OTR density was significantly higher in the withdrawn group than the no hormone group (p = 0.0002) and the sustained group (p = 0.0033, Figure 3E).

**Figure 3:**
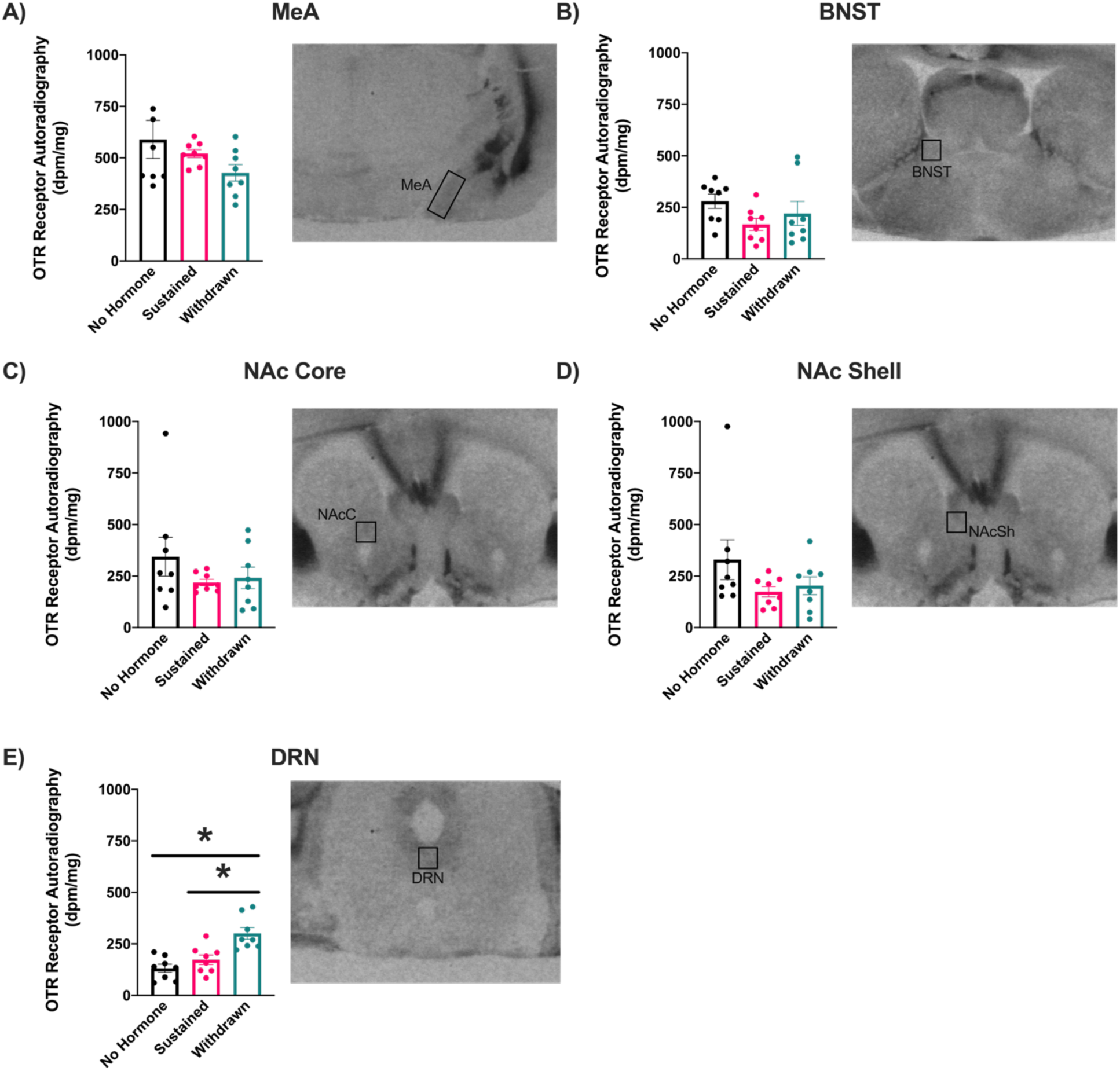
Effect of Estrogen-Withdrawal on OTR density in estrogen-sensitive PVH efferents. OTR autoradiographic binding did not differ across hormone conditions in A) the MeA, B) the BNST, C), the NAc Core, or D) the NAc Shell. E) In the DRN, however, withdrawn females had significantly higher OTR autoradiographic binding than sustained or no hormone controls. Representative autoradiographs with analysis boxes over the region of interest are shown next to each data set. Data presented as mean ± SEM, * *p* < 0.05.

### Experiment 4: Effects of blocking oxytocin receptors in the DRN on anxiety-like behaviors

In experiments 1-3, we found that estrogen withdrawal following HSP increased the number of OT-ir cells in the PVH and the density of OTR in the DRN. These changes were accompanied by an increase in anxiety-like behaviors in the elevated plus and open field. We therefore hypothesized that increased oxytocin signaling between the PVH and DRN results in a higher anxiety phenotype during the postpartum period. To test this hypothesis, we used a behavioral pharmacology approach to block OTRs in the DRN following HSP and measured the impact on anxiety-like behaviors.

#### Cannulae Placement

One hamster did not recover from cannulation surgery and the cannulae on eight hamsters did not remain intact and/or patent throughout the HSP protocol. In the remaining 51 hamsters, histological verification of cannulae placement using post-mortem ink injections showed that drug injections targeted the caudal DRN (5.4 - 5.7 mm posterior to Bregma, Figure 4A, (30)) in 32 subjects. Subjects with cannulae outside of the DRN were misplaced either in the lateral periaqueductal gray (LPAG; *n* = 5), the fourth ventricle (*n* = 6), the lateral dorsal tegmental nucleus (n = 5) or, more rarely, the medial longitudinal fasciculus (n = 3); these animals were considered misses and removed from the analyses.

**Figure 4:**
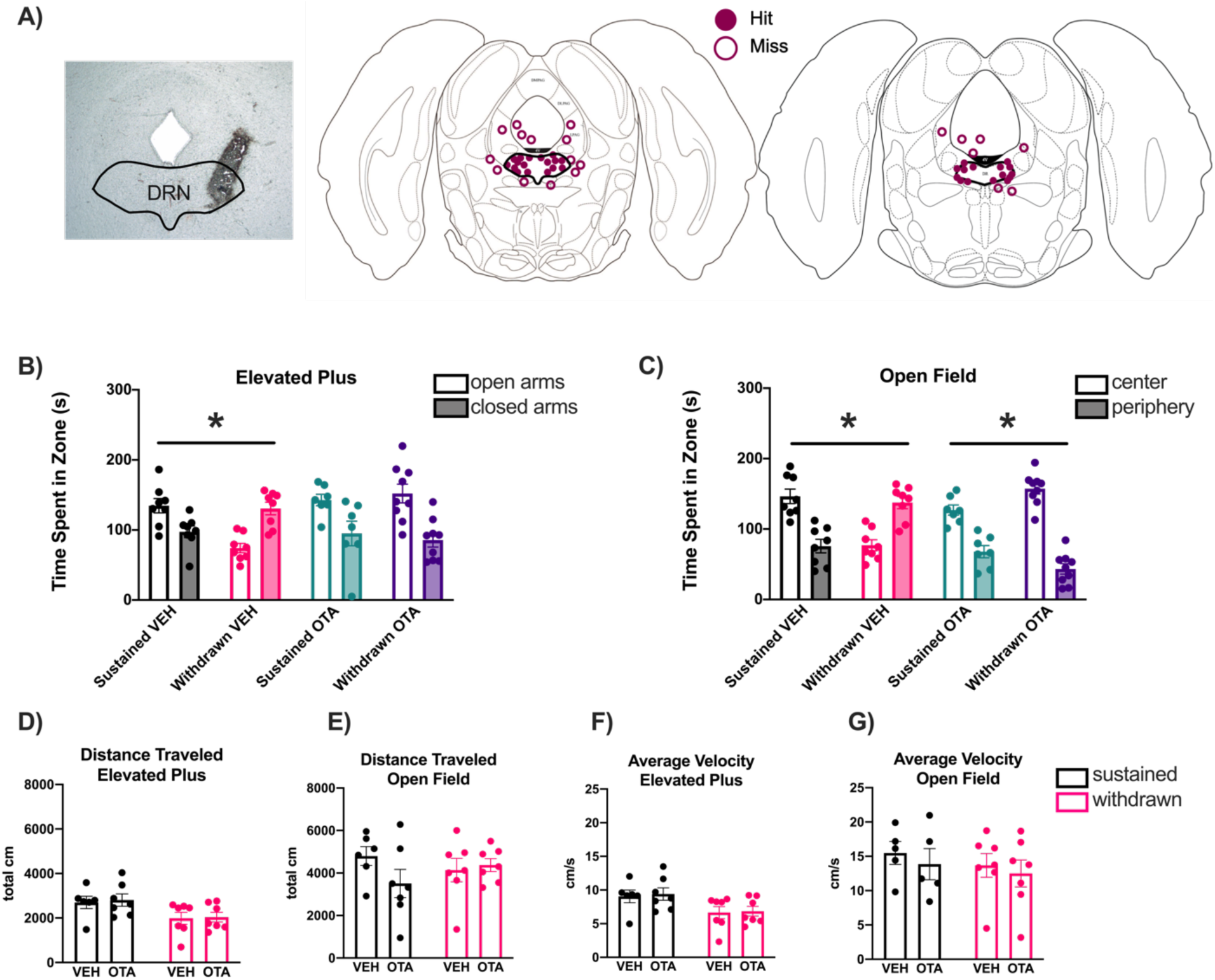
Effect of Blocking OTRs in DRN during Estrogen-Withdrawal on Anxiety-Like Behaviors. A) Post-mortem localization of ink injections was used to determine whether placements accurately targeted the DRN (“hit,” filled circles) or were misplaced (“miss,” open circles) and therefore eliminated from analyses. OTA injection into the DRN reverses the high-anxiety behavioral phenotype seen in withdrawn females in B) the elevated plus and C) the open field. Although there was a main effect of hormone condition on distance traveled in the elevated plus, differences in anxiety-like behavior are unlikely to be mediated by a more general locomotor deficit, as there was no difference in distance traveled (D and E) or average velocity (F and G) between sustained and withdrawn females who received vehicle (VEH) injections or OTA injections. Data presented as mean ± SEM, * *p* < 0.05.

#### Elevated Plus

There was a significant interaction between hormone condition (sustained vs. withdrawn) and drug condition (OTA vs. vehicle) (F(1,28) = 8.916, p = 0.0058) in the elevated plus (Figure 4B). Post hoc tests revealed that in females who received vehicle injections into the DRN, the sustained and withdrawn groups differed from each other (t(28) = 3.532, p = 0.0029): sustained animals who received vehicle spent more time in the open arms of the elevated plus, whereas withdrawn animals who received vehicle spent more time in the closed arms of the elevated plus. In females who received OTA injections into the DRN, the sustained and withdrawn groups did not differ from each other (t(28) = 0.7018, p = 0.9772); both groups spent more time in the open arms of the elevated plus. Together, these data suggest that postpartum estrogen withdrawal increases anxiety-like behavior in the elevated plus, and that this increase can be prevented by blocking OT receptors in the DRN.

#### Open Field

There was a significant interaction between hormone condition and drug condition (F(1,28) = 44.73, p < 0.0001) in the open field (Figure 4C). Post hoc tests revealed that in females who received vehicle injection into the DRN, the sustained and withdrawn groups differed from each other (t(28) = 6.701, p < 0.0001): sustained animals who received vehicle spent more time in the center of the open field, whereas withdrawn animals who received vehicle spent more time in the periphery of the open field. In females who received OTA injections into the DRN, the sustained and withdrawn groups also differed from each other (t(28) = 2.773, p = 0.0195): although both sustained and withdrawn females spent more time in the center of the open field, withdrawn females who received OTA spent more time in the center than did sustained females who received OTA. Together, these data suggest that postpartum estrogen withdrawal increases anxiety-like behavior in the open field, and that this increase can be prevented by blocking OT receptors in the DRN.

#### Locomotor Behavior

To assess whether the significant interactions between hormone condition and drug condition in the elevated plus and open field could be explained by broader locomotor deficits, distance traveled (total cm) and average velocity (cm/s) were measured in both apparatuses. There was no significant interaction between hormone and drug condition on distance traveled in the elevated plus (F(1,23) = 0.01153, p = 0.9154, Figure 4D) or the open field (F(1,23) = 2.208, p = 0.1509, Figure 4E). Interestingly, there was a main effect of hormone condition on distance traveled in the elevated plus (F(1,23) = 8.002, p = 0.0095), but post hoc tests did not detect significant differences between sustained and withdrawn females who received vehicle injections (t(23)=1.887, p = 0.1437) or OTA injections (t(23)=2.119, p = 0.0902). There was no significant interaction between hormone and drug condition on average velocity in the elevated plus (F(1, 23) = 0.008757, p = 0.9263, Figure 4F) or the open field (F(1,20) = 0.01354, p = 0.9085, Figure 4G). This suggests that group differences in the elevated plus and open field in Experiment 4 are reflective of differences in anxiety-like behaviors rather than nonspecific effect of drug or hormone condition on locomotion.

## Discussion

Here we demonstrate for the first time that estrogen withdrawal following HSP increases behavioral measures of anxiety in Syrian hamsters. Further, these increases in anxiety-like behavior are concurrent with increases in OT-producing cells in the PVH and increased OTR density in the DRN. Finally, blocking OTRs in the DRN during estrogen withdrawal prevents the increase in anxiety-like behavior in withdrawn females, but has minimal effect in females who continued to receive estrogen throughout the simulated postpartum period. Together, these results suggest a model in which postpartum estrogen withdrawal increases oxytocin signaling in the PVH and DRN, either independently or as part of a neural pathway, to increase anxiety-like behavior during the postpartum period.

These data add to a growing body of literature that uses HSP to model the impact of postpartum estrogen withdrawal, and expands the model to Syrian hamsters and to oxytocinergic neural circuitry. Although the HSP model does not provide a comprehensive simulation of all the hormonal changes that occur during the peripartum period, it allows researchers to isolate and manipulate peripartum estrogen fluctuations as a single independent variable, permitting causal inferences about resulting changes in the brain and behavior. Further, unlike in women, most common laboratory rodents experience postpartum and lactational estrus (31), during which estrogen levels rise sharply; in this way, HSP better models the human experience of postpartum estrogen withdrawal than would naturally-parturient rodents.

Differences in model organisms may influence neural and behavioral outcomes of HSP. Similar to in the current study, estrogen withdrawal in Sprague-Dawley rats (11) and ICR mice (14,15) increases behavioral indicators of anxiety, suggesting that elevated anxiety following postpartum estrogen withdrawal is a robust behavioral phenomenon and may reflect an adaptive response that serves to increase vigilance during a vulnerable time. In contrast, the current study did not find an effect of estrogen withdrawal on sucrose preference, despite this being reported in rats (8,12). This may reflect differences in the behavioral ecology of the two species. Although hamsters are capable of forming sucrose preferences (32), they consume less water per body weight than rats (33), possibly due to their native desert habitat. As such, a drinking-based assay may not be the most robust test of anhedonia in this species. Future studies may wish to use other assays, such as intracranial self-stimulation (9), to test the effect of estrogen withdrawal on anhedonia in hamsters.

Although the HSP model has low ecological validity compared to naturally-parturient rodents who can engage in maternal and lactational behaviors, the current findings are corroborated by data from postpartum rats (34,35) and rabbits (36), in which OT-immunoreactivity is increased in the PVH. Caldwell et al. also found increased OT-immunoreactivity in the SON of postpartum female rats, which differs from our findings using the HSP model; this suggests that while OT levels in the PVH and SON are both dynamic during the peripartum period, only PVH OT levels are likely to be mediated by estrogen levels. Of interest, a recent study that used a gestational stress model of postpartum depression found that postpartum rats who had undergone restraint stress during pregnancy had decreased OT immunoreactivity and OT mRNA expression in the PVH (37). On the surface, this conflicts with our finding that OT immunoreactivity and OT mRNA levels are increased following estrogen withdrawal; however, the gestational stress study only compared postpartum females with and without gestational stress, so it is unclear whether postpartum OT levels are higher or lower relative to pregnant, cycling, or ovariectomized females. Future experiments may benefit from combining the HSP model with restraint stress to tease apart the effects of estrogen withdrawal and gestational stress on OT in the PVH.

Our finding that OTR autoradiographic binding is increased in the DRN of estrogen withdrawn females is also supported by recent data from naturally-parturient females showing that OTR autoradiographic binding and OT-ir fibers are increased in the DRN of postpartum rats (38). What’s more, viral-mediated knockdown of OTR mRNA in the DRN of postpartum rats decreases anxiety-like behavior in the elevated plus (38), mirroring our finding that pharmacological blockade of OTRs in the DRN decreased anxiety-like behavior in the elevated plus during estrogen withdrawal. When coupled with previous data, this suggests that 1) oxytocin in the DRN has anxiogenic effects on behavior and 2) these effects may depend on postpartum estrogen withdrawal. More broadly, these data join a growing body of literature challenging the dogma that OT is primarily anxiolytic (18,39) and suggests the need for more site-specific investigations of OT’s role in modulating affective behaviors.

The mechanism by which estrogen withdrawal increases OT-ir cells in the PVH is not known. The current experiments suggest that it is not an artifact of decreased OT release, as OT mRNA levels also increase substantially in withdrawn females, which would not be expected with decreased release. Further, it is unlikely to reflect an increase in the total number of cells, as we did not observe an increase in NeuN-immunoreactivity between hormone conditions. One intriguing possibility is that postpartum estrogen withdrawal is causing PVH cells to switch their neurochemical phenotype, either temporarily or permanently. This is not without precedent in the hypothalamus: in pregnant and lactating mice, tuberoinfundibular neurons in the arcuate nucleus begin synthesizing and releasing met-enkephalin rather than dopamine (40). The mechanism of this phenotypic switch depends on elevated prolactin levels during pregnancy suppressing dopamine synthesis in an opiate-dependent manner. It is possible that the massive fluctuations in estrogen during pregnancy and the postpartum period may similarly result in neurochemical plasticity at the level of individual neurons in order to facilitate adaptive behavioral changes during the peripartum period.

We propose that estrogen withdrawal increases OT signaling between the PVH and DRN in order to increase anxiety during the postpartum period (Figure 5). The PVH sends moderately dense projections to the DRN, terminating primarily in the A10dc dopaminergic cell group (41). These dopaminergic DRN cells have been associated with arousal (42), and project to nuclei in the extended amygdala that are known to regulate anxiety (43,44). Recent work by Grieb et al (2019) demonstrates that approximately one third of these dopamine cells express OTRs, and the number and percentage of dopamine cells expressing OTRs were higher in postpartum rats than in diestrus virgin rats (38). In addition to dopaminergic cells, the raphe nuclei, including the DRN, are the primary source of forebrain-projecting serotonin in the brain (45). Approximately half of raphe serotonin neurons contain OTRs, which modulate serotonin release to regulate anxiety (46). Given these neuroanatomical and functional substrates, it seems likely that estrogen withdrawal increases OT signaling between the PVH and the DRN in order to increase anxiety during the postpartum period, either by acting directly at dopaminergic neurons or indirectly at serotonin neurons. Moving forward, experiments using chemogenetic inhibition of the PVH-DRN pathway are necessary to demonstrate its requirement for estrogen withdrawal-dependent increases in anxiety in the postpartum period. Further, determining whether dopamine and/or serotonin signaling is altered in the DRN following estrogen withdrawal will be useful in providing a more complete understanding of how this circuit may be altered during the postpartum period. Ultimately, these data suggest a novel neural pathway in which peripartum estrogen fluctuations lead to neural plasticity that impacts postpartum affective behaviors.

**Figure 5:**
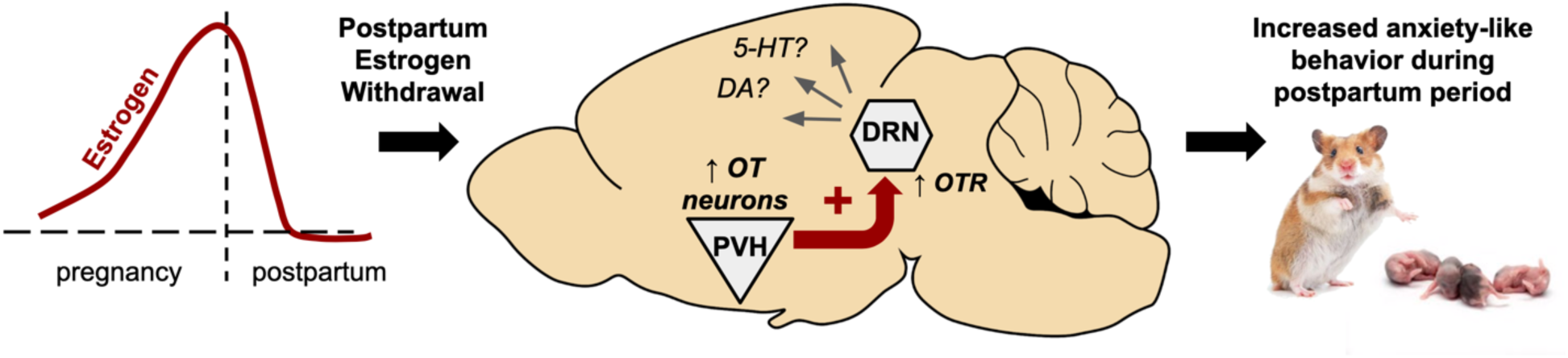
Proposed Model. Based on our data, we propose a model in which postpartum estrogen withdrawal increases oxytocin signaling between the PVH and DRN to increase anxiety-like behavior during the postpartum period. Specifically, we suggest that estrogen withdrawal leads to an increase in OT neurons in the PVH and a simultaneous increase in OTR density in the DRN, indicating a pathway-specific increase in OT neurotransmission. OT from the PVH may act directly on dopamine (DA) neurons in the DRN to increases baseline arousal, or indirectly on serotonin (5-HT) neurons in the DRN to increase anxiety, during the early postpartum period.

## Acknowledgements

The authors thank Haley Bianco and Taylor Friendt for their assistance with data collection, and Nicholas Jones and Erin Haughee for their management of the Haverford College Vivarium. This work was supported in part by a Haverford College faculty research grant (LEB) and NIH grant R01MH110212 (HEA). Portions of this work were presented previously at the Society for Neuroscience and Society for Behavioral Neuroendocrinology annual meetings. This manuscript was submitted simultaneously to BiorXiv.org as a preprint.

