## Supplementary Figure 1 for "Estrogen withdrawal alters oxytocin signaling in the paraventricular hypothalamus and dorsal raphe nucleus to increase postpartum anxiety"

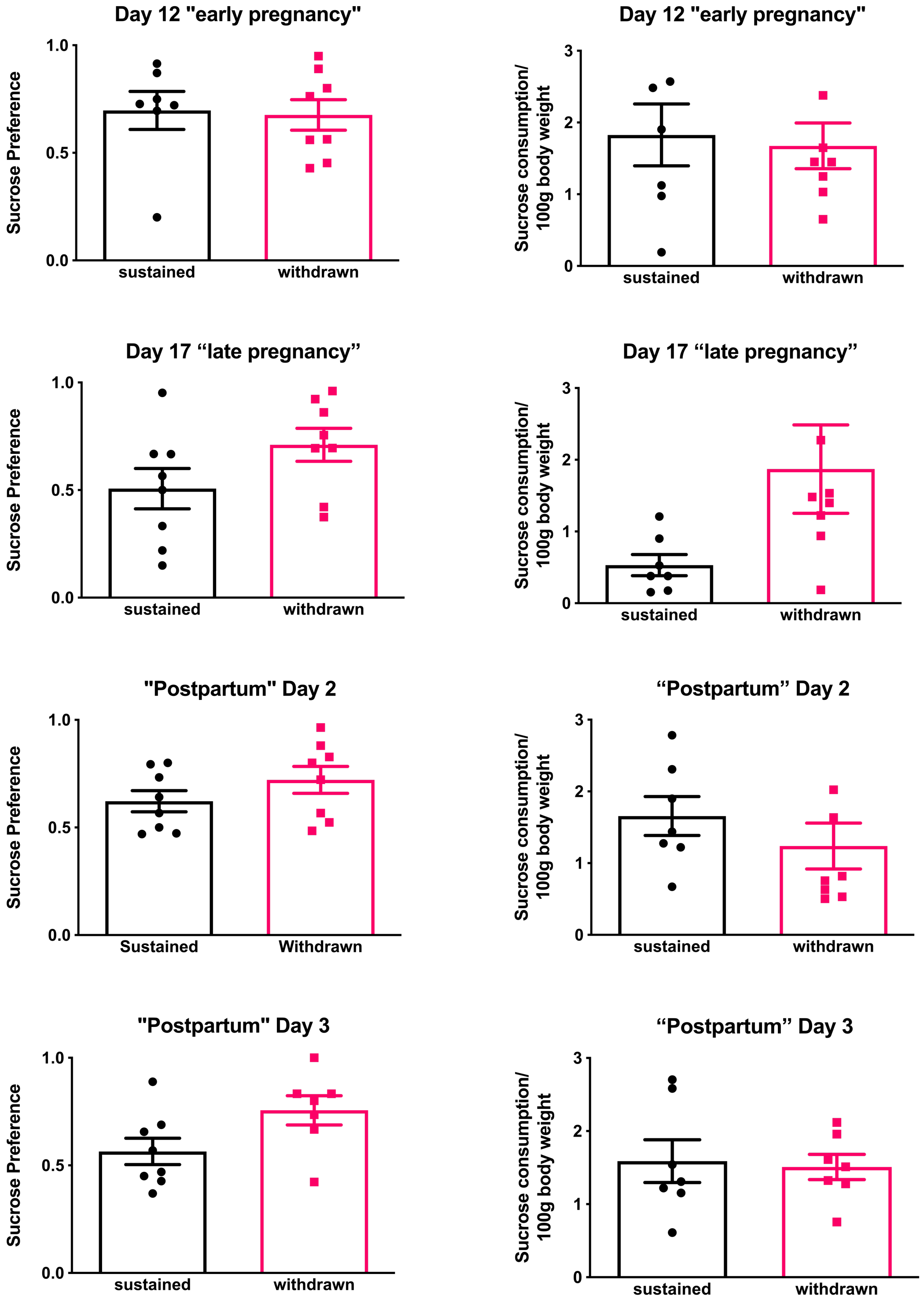


**Supplementary Figure 1: Effect of Estrogen-Withdrawal on Sucrose Preference.** There was no effect of hormone treatment on sucrose preference or sucrose consumption on any day. Data presented as mean ± SEM.
